# Identification of Wee1 as Target in Combination with Avapritinib for the Treatment of Gastrointestinal Stromal Tumor

**DOI:** 10.1101/2020.06.07.138693

**Authors:** Shuai Ye, Dinara Sharipova, Marya Kozinova, Lilli Klug, Jimson D’Souza, Martin G. Belinsky, Katherine J. Johnson, Margret B. Einarson, Karthik Devarajan, Yan Zhou, Samuel Litwin, Michael C. Heinrich, Ronald DeMatteo, Margaret von Mehren, James S. Duncan, Lori Rink

## Abstract

Management of gastrointestinal stromal tumor (GIST) has been revolutionized by the identification of activating mutations in KIT and PDGFRA, and the clinical application of receptor tyrosine kinase (RTK) inhibitors in the advanced disease setting. Stratification of GIST into molecularly defined subsets provides insight into clinical behavior and response to approved targeted therapies. Although these RTK inhibitors are effective in the majority of GIST, resistance to these agents remains a significant clinical problem. Development of effective treatment strategies for refractory GIST subtypes requires identification of novel targets to provide additional therapeutic options. Global kinome profiling has the potential to identify critical signaling networks and reveal protein kinases that are essential in GIST. Using Multiplexed Inhibitor Beads and Mass Spectrometry, we explored the majority of the kinome in GIST specimens from the three most common molecular subtypes to identify novel kinase targets. Kinome profiling revealed distinct signatures in GIST subtypes and identified kinases that are universally activated in all GIST, as well as kinases that are unique to each subtype. Kinome profiling in combination with loss-of-function assays identified a significant role for the G2-M tyrosine kinase, Wee1, in GIST cell survival. *In vitro* and *in vivo* studies revealed significant efficacy of MK-1775 (Wee1 inhibitor) in combination with avapritinib in *KIT* and *PDGFRA*-mutant GIST cell lines, and notable efficacy of MK-1775 as a single agent in the *PDGFRA*-mutant line. These studies provide strong preclinical justification for the use of MK-1775 in GIST.

## Introduction

Gastrointestinal Stromal Tumors (GIST) are the most common mesenchymal tumors of the gastrointestinal tract, with 5000-6000 new cases diagnosed in the US annually (1). These tumors are characterized by near-universal expression of the receptor tyrosine kinase (RTK) KIT, and the majority of GIST harbor constitutively active mutant isoforms of KIT (70-80%) or the related RTK, PDGFRA (5-7%) (2). The ∼10-15% of GIST that lack mutations in these genes often exhibit genetic or epigenetic deficiencies in the succinate dehydrogenase (SDH) complex of the respiratory chain (3, 4), and are referred to as SDH-deficient (SDH-d). Therapeutic targeting of GIST with the front-line RTK inhibitor, imatinib mesylate (IM) along with three other FDA approved agents (sunitinib, regorafenib and repretinib) has transformed therapy for advanced, unresectable GIST. However, this “one size fits all” approach to GIST treatment fails to address the molecular and clinical heterogeneity of these tumors. Tumor genotype has been shown to be an independent prognostic factor and a predictor of IM response in GIST (5). The majority of GIST harbor mutations in *KIT* that affect the juxtamembrane domain encoded by exon 11. While tumors with mutations in this region generally respond well initially to IM therapy, they may exhibit negative prognostic features and aggressive biology (6). In contrast, *PDGFRA-*mutant and SDH-d GIST may exhibit a more indolent clinical course (7), however these GIST subtypes demonstrate little or no response to IM (8, 9) or to other approved therapies. The most common *PDGFRA* mutation found in GIST, the D842V substitution, is particularly insensitive to IM. Avapritinib (BLU-285, Blueprint Medicines), a highly selective inhibitor of *KIT* exon 17 and *PDGFRA* exon 18 activation loop mutants, has demonstrated efficacy *in vitro* (10) and *in vivo* (11). Phase I testing (NAVIGATOR study, NCT02508532) has demonstrated notable efficacy for exon 18 *PDGFRA*-mutant GIST (12), leading to FDA approval for the use of avapritinib in unresectable or metastatic *PDGFRA* exon 18-mutant GIST in January 2020.

Although these RTK inhibitors are effective in most GIST, primary and acquired resistance remains a significant clinical obstacle. For clinical management of refractory GIST to improve, new therapeutic targets must be identified. Finding a better way forward will require a more complete understanding of how the particular molecular aberrations in GIST subsets affect tumor signaling pathways and ultimately impact clinical behavior and therapeutic response. Differences in global gene expression and genomic profiles have been reported for GIST subtypes (3, 13, 14), however kinome profiling of GIST has not been performed to date. Global kinome profiling has the potential to identify essential signaling networks and reveal protein kinases that are critical in GIST. Protein kinases are highly druggable, with more than 45 FDA approved kinase inhibitors (15), the majority of which are used clinically to treat malignancies. Several chemical proteomics approaches have been developed that measure levels of a significant proportion of the kinome in cells and tissues, including Kinobeads, Kinativ and Multiplexed Inhibitor Beads and Mass Spectrometry (MIB-MS) (16-18). MIBs consists of a layered mixture of immobilized ATP-competitive pan-kinase inhibitors that enriches endogenous protein kinases from cell lysates based on affinity of individual kinases for the different immobilized inhibitors, their kinase abundance, and/or kinase activation state (17).

In this work, using MIB-MS (19, 20), we explored the majority of the kinome in treatment naïve primary GIST specimens from three GIST subtypes (*KIT*-mutant, *PDGFRA*-mutant and SDH-d GIST) in order to identify potential novel targets. Using this proteomics approach, we demonstrated that the three GIST subtypes have distinct kinome profiles, and identified kinases that are universally activated in all GIST, as well as kinases that are unique to each subtype. Finally, kinome profiling in combination with loss-of-function validation assays revealed a significant role for the G2-M tyrosine kinase, Wee1, in GIST survival. We also report significant efficacy of MK-1775 (Wee1 inhibitor) as a monotherapy, and in combination with avapritinib in a novel GIST cell line driven by an activating *PDGFRA* D842V mutation. The combination was also effective in controlling the growth of these PDGFRA-driven GIST cells in 3-dimensional spheroid culture. Furthermore, dual inhibition of Wee1 and KIT/PDGFRA in GIST xenografts provided impressive, extended disease stabilization and improved survival.

## Results

### Kinome profiling of primary GIST using MIB-MS

To explore the kinome landscapes among the three molecular subtypes of GIST, we performed MIB-MS profiling on 33 IM-naïve primary gastric GIST specimens which included the following subtypes: 1) *KIT* exon 11 mutants (N=15), 2) *PDGFRA* mutants (N=10) and 3) *KIT/PDGFRA*-wild-type (WT) GIST (N=8) (**Table 1)**. The *KIT/PDGFRA*-WT GISTs include 7 SDH-d and one GIST that lacked *SDH* mutations and was shown to be SDH intact by SDHB immunohistochemistry (21). We also kinome profiled 9 normal gastric tissues from donors without a history of kinase inhibitor therapy. To quantify the MIB-bound kinome of GIST tissues, we performed label-free protein quantitation (LFQ) using the MaxLFQ algorithm (22) in combination with a super-SILAC (s-SILAC) (23) internal standard to control for variations in kinase MIB-binding and/or LC-MS/MS retention time reproducibility (**Figure 1A)**. In total, we measured MIB-binding values for 296 kinases across these GIST samples, with 242 kinases quantitated in >70% of tissues profiled and 156 kinases measured in every MIB-MS run. The average number of kinases measured for each sample was 254 (**Figures 1B-C**, and **Data file S1**). Principal component analysis (PCA) of MIB-MS profiles revealed that the GIST kinome is overall distinct from normal gastric tissues (**Figure 2A**). Furthermore, PCA of the kinome profiles revealed *KIT-* and *PDGFRA*-mutant GIST grouped distinctly from KIT/PDGFRA-WT GIST (**Figure 2B-C**). One exception to this was *KIT/PDGFRA*-WT33 sample that clustered closer to *KIT*-mutant GIST samples. Interestingly, this sample was distinct from other KIT/PDGFRA-WT GIST samples in that it possessed an intact SDH complex, and most likely has an unknown driver mutation.

**Table 1:** Genotype information for GIST patient samples

| Table 1: Genotype information for GIST patient samples |  |  |
| --- | --- | --- |
| Sample number | Group | Mutation(s)<sup>a</sup> |
| 1 | KIT mutant | *KIT* exon 11: p.Val559Ala |
| 2 | KIT mutant | *KIT* exon 11: p.Val559Asp |
| 3 | KIT mutant | *KIT* exon 11: p.Val560Asp |
| 4 | KIT mutant | *KIT* exon 11: p.Val559Pro |
| 5 | KIT mutant | *KIT* exon 11: p.Val560Asp |
| 6 | KIT mutant | *KIT* exon 11: p.Val560Asp |
| 7 | KIT mutant | *KIT* exon 11: p.Trp557Arg |
| 8 | KIT mutant | *KIT* exon 11: p.Val559Gly |
| 9 | KIT mutant | *KIT* exon 11: p.Trp557Gly |
| 10 | KIT mutant | *KIT* exon 11: p.Val560Asp |
| 11 | KIT mutant | *KIT* exon 11: p.Val559Ala |
| 12 | KIT mutant | *KIT* exon 11: p.Trp557_Val559delinsPhe |
| 13 | KIT mutant | *KIT* exon 11 indel |
| 14 | KIT mutant | *KIT* exon 11: p.Trp557_Val559delinsCys |
| 15 | KIT mutant | *KIT* exon 11: p.Pro573_Glu583dup |
| 16 | PDGFRA mutant | *PDGFRA* exon 14: p.Asn659Tyr; exon 18: p.Tyr849Cys |
| 17 | PDGFRA mutant | *PDGFRA* exon 18: p.Asp842Val |
| 18 | PDGFRA mutant | *PDGFRA* exon 18: p.Asp842Val |
| 19 | PDGFRA mutant | *PDGFRA* exon 18: p.Asp842Ile |
| 20 | PDGFRA mutant | *PDGFRA* exon 12: p.Val561Asp |
| 21 | PDGFRA mutant | *PDGFRA* exon 18: p.Asp842Val |
| 22 | PDGFRA mutant | *PDGFRA* exon 18: p.Asn842Lys |
| 23 | PDGFRA mutant | *PDGFRA* exon 12: p.Val561Asp |
| 24 | PDGFRA mutant | *PDGFRA* exon 18: p.Asp842Val |
| 25 | PDGFRA mutant | *PDGFRA* exon 18: p.Asp842Val |
| 26 | KIT/PDGFRA wild type | *SDHC* exon 4: p.Gly75Asp |
| 27 | KIT/PDGFRA wild type | none identified |
| 28 | KIT/PDGFRA wild type | none identified |
| 29 | KIT/PDGFRA wild type | none identified |
| 30 | KIT/PDGFRA wild type | *SDHA* exon 14: p.Tyr629Phe; exon 15: p.Val657Ile |
| 31 | KIT/PDGFRA wild type | *SDHA* exon 7: p.Thr273Ile; exon 10: p.Gly453Arg |
| 32 | KIT/PDGFRA wild type | none identified |
| 33<sup>b</sup> | KIT/PDGFRA wild type | none identified |

<sup>a</sup>Amino acid numbering from isoforms NP\_000213.1 (KIT), NP\_006197.1 (PDGFRA), NP\_004159.2 (SDHA), NP\_002992.1 (SDHC)
<sup>b</sup>SDH-intact WT

**Figure 1.**
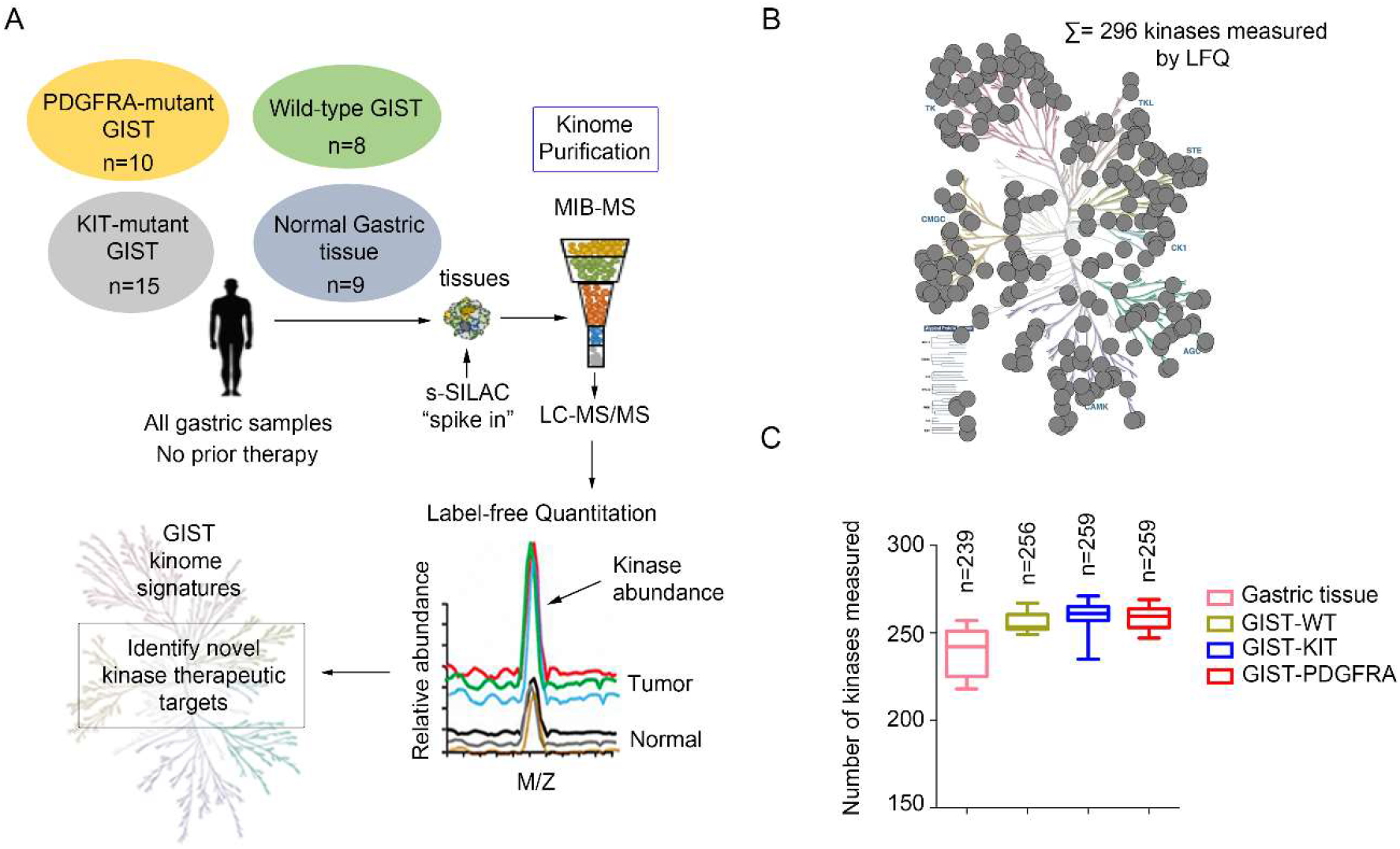
Characterizing the GIST kinome in primary tumors using MIB-MS to identify novel therapeutic targets. **(A)** Schematic of experimental approach. MIB-MS was used to quantify the kinase abundance in GIST patient tumors (untreated, gastric primary GIST from three molecular subtypes: *KIT*-mutant, n=15; *PDGFRA*-mutant, n=10; wild-type GIST, n=8, and normal gastric tissue, n=9) to map the proteomic landscape of the kinome and identify novel targets. Kinase levels in tissues were determined using a combination of Label-Free Quantitation (LFQ) and super-SILAC (s-SILAC). **(B)** Kinome tree depicts fraction of kinome quantitated by MIB-MS and frequency across 42 samples measured. **(C)** Average number of kinases detected by MIB-MS profiling broken down by tissue type.

**Figure 2.**
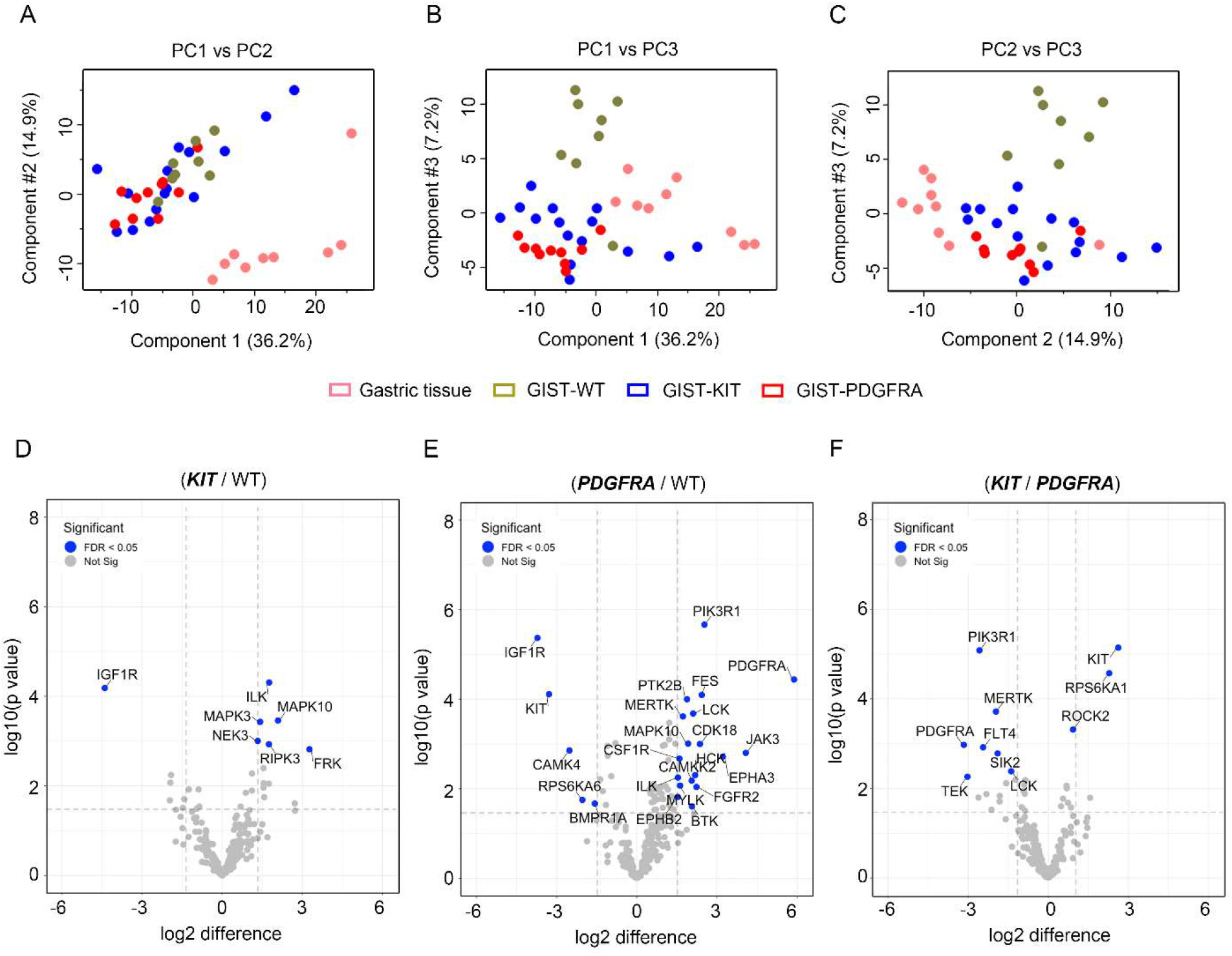
Mapping the distinct kinome signatures among GIST subtypes. **(A-C)** Principal Component Analysis, including PC1 versus PC2 (A), PC1 versus PC3 (B), and PC2 versus PC3 (C) of MIB-MS in three GIST subtypes (*KIT*-mutant (blue), *PDGFRA*-mutant (red), wild-type (green) and normal gastric tissue (pink). **(D)** Volcano plot comparisons of KIT-mutant versus WT, *PDGFRA*-mutant versus WT, and **(F)** *KIT*-mutant versus *PDGFRA*-mutant GIST MIB-MS kinome profiles. Differences in kinase log2 LFQ intensities among tumors and normal tissues determined by paired *t*-test Benjamini-Hochberg adjusted *P* values at FDR of <0.05 using Perseus software.

### Mapping the distinct kinome signatures among GIST subtypes

Volcano plot analysis of kinase log_2_ LFQ values (**Figures 2D-F**) revealed kinases exhibited differential protein abundance among GIST subtypes (**Data file S1**). A scatterplot comparing LFQ or s-SILAC-determined log2 differences in kinases levels among tissues shows the two quantitative methods displayed substantial overlap, validating the majority of kinase measurements (**Figure S1A-F**). Some kinases were not quantitated by the super-SILAC method, which can be attributed to low or absent expression of kinase in SILAC-labeled cell line cocktail. IGF1R was the top ranking kinase elevated in *KIT/PDGFRA*-WT versus *KIT*- or *PDGFRA*-mutant subtypes, as previously described (24, 25) **(Figures 2D-E and S1B-D)**. *PDGFRA*-mutant tumor kinome profiles greatly differed from *KIT/PDGFRA*-WT tumors, with many kinases showing increased protein levels in PDGFRA-mutant tumors relative to wild-type (**Figure 2E and S1D**). Elevated levels of several RTKs, including PDGFRA, MERTK, EPHB2, EPHA3, CSF1R, and FGFR2 were observed in *PDGFRA*-mutant tumors vs WT tumors, as well as increased levels of PIK3R1, the regulatory subunit of PIK3CA, which has been shown to be an an important downstream signaling effector of both KIT and PDGFRA (26) (**Figure 2E, S2C-D**). Notably, many of the elevated kinases in the *PDGFRA*-mutant tumors were kinases related to immune cell function, including HCK, LCK, BTK, CSF1R and MERTK. These findings are consistent with a recent report from Vitiello et al. (27) demonstrating increased immune cells present in *PDGFRA*-mutant GIST. Conversely, increased ROCK2 was detected in *KIT*-mutant vs. *PDGFRA*-mutant tumors (**Figure 2F, S2F)**. ROCK2 has been associated with increased aggressiveness and metastasis, as well as poor overall survival in several malignancies (28).

### Targeting the GIST tumor kinome signature identifies WEE1 as candidate target

Next, we explored kinases commonly overexpressed among *KIT*- and *PDGFRA*-mutant tumors relative to normal gastric tissue, with the goal to identify new kinase targets to exploit in GIST. As expected, volcano plot analysis showed significant differences between normal gastric tissue and GIST-mutant tumors, with numerous kinases expressed at higher levels in *KIT*- and *PDGFRA*-mutant tumors relative to normal gastric tissues (**Figure 3A, and Data file S1**). The majority of LFQ-determined kinase measurements were confirmed by super-SILAC, representing high-confidence kinase signatures (**Figure 3B**). Significant elevation of KIT, PRKCQ and FGFR1, all of which have been previously shown to be up-regulated in GIST (29, 30), as well as kinases associated with regulation of cell cycle (WEE1 and CDK4), NF-kB signaling (TBK1) and stress response signaling (MAP3K3, STK3, MAPK10 and PRKD1) (**Figure 3C-D**), was seen. To explore functional relevance of the high-confidence kinases commonly elevated in *KIT*- and *PDGFRA*-mutant tumors, we designed a kinase-centric siRNA library to identify kinases that are critical for *KIT-* and *PDGFRA*-mutant GIST cell survival. This siRNA library contained pooled siRNAs targeting each of the 13 kinases identified in the kinome profiling experiment. Synthetic lethal screens were performed using an isogenic pair of cell lines: GIST-T1+Cas9 (KIT driven) and GIST-T1+D842V KIT^KO^ (PDGFRA D842V driven). Positive controls for the screen included siKIT (GIST-T1+Cas9) and siPDGFRA (GIST-T1+D842V KIT^KO^), while siGL2 served as negative control for both lines. Knockdown of the majority of the kinases in the screen showed minimal impact on cell viability (**Figure 3E**). However, siRNA-mediated depletion of WEE1 led to significantly decreased viability in both isogenic lines (viability score = 0.48 in GIST-T1+Cas9; 0.41 in GIST-T1+D842V KIT^KO^), while knockdown of MAP3K3 significantly affected viability in the *PDGFRA*-mutant cell line (viability score = 0.39). Viability reductions approaching that of the KIT and PDGFRA positive controls were seen for these two kinases. Greater than seventy percent Wee1 knockdown was achieved in both cell lines (**Figure 3F)**. Interestingly, depletion of MAP3K3, known to promote ovarian and NSCLC tumor growth (31, 32), inhibited *PDGFRA*-mutant GIST cell viability, however no selective small molecule inhibitors are currently available to explore targeting MAP3K3 in GIST. Overexpression of Wee1 has also been observed in numerous malignancies, including breast and melanoma (33). MK-1775 (adavosertib, AZD1775), a selective inhibitor targeting Wee1, is under investigation in clinical trials, and recently several preclinical studies have suggested significant synergy when Wee1 inhibitors was combined with other kinase inhibitors, including the mTOR inhibitor, TAK228 (34) and the AURKA inhibitor, alisertib (35). Our GIST kinome profiling data along with these initial cell viability studies suggests that Wee1 could be a plausible drug target in mutant GIST, either alone or in combination with existing therapies.

**Figure 3.**
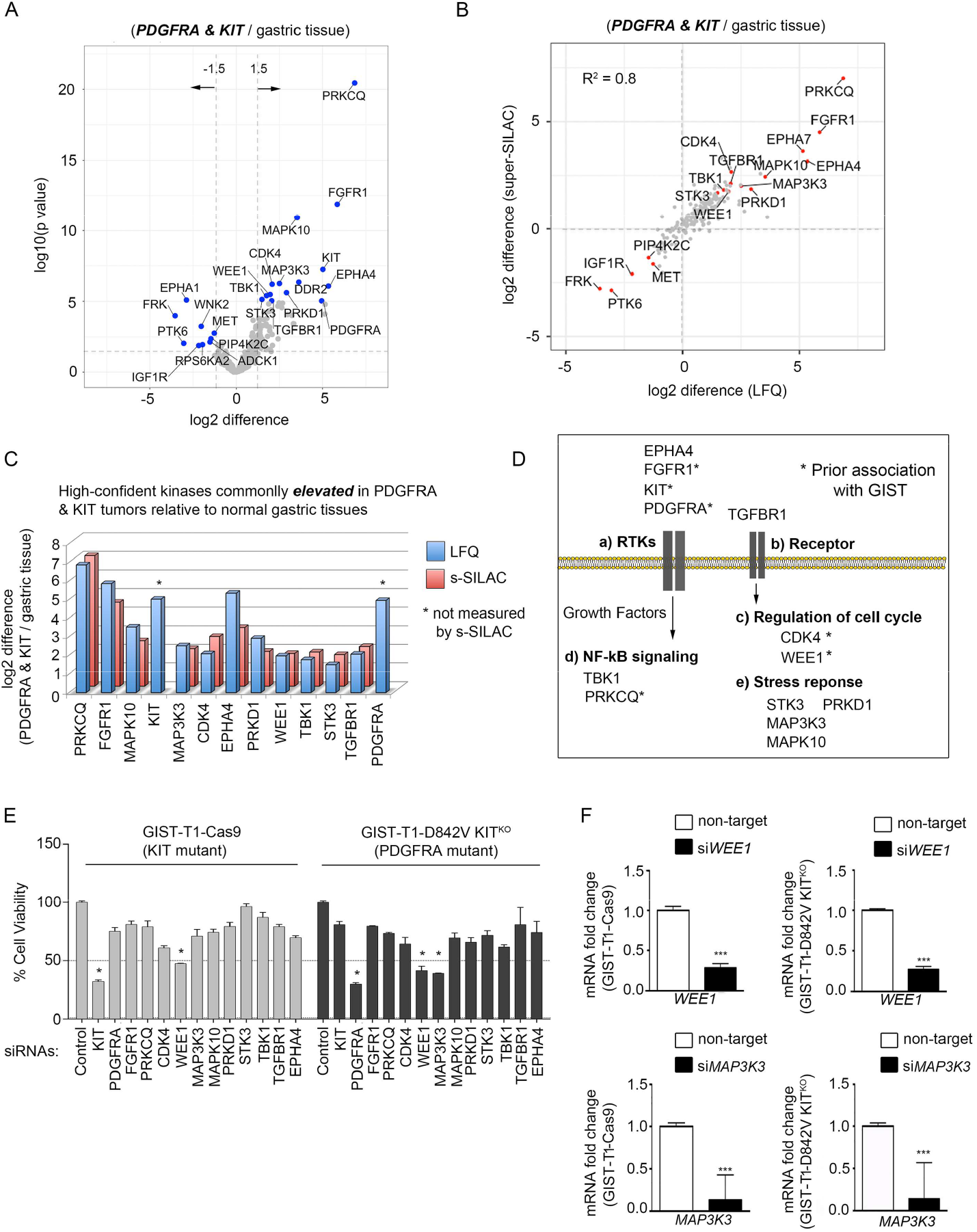
Targeting the mutant-GIST kinome signature identifies WEE1 as candidate target. **(A)** Volcano plot comparisons of *KIT-* and *PDGFRA-*mutant GIST vs. normal gastric tissue MIB-MS kinome profiles. Differences in kinase log2 LFQ intensities among tumors and normal tissues determined by paired *t*-test Benjamini-Hochberg adjusted *P* values at FDR<0.05 using Perseus Software. (**B**) Scatterplot depicts overlap in kinases elevated or reduced determined by LFQ or s-SILAC. Regression analysis (R^2^) among quantitative methods was performed in Perseus Software. Differential expressed kinases commonly identified by LFQ and s-SILAC quantitation (FDR<0.05) are labeled. **(C)** Bar graph depicts high-confident kinases log2 LFQ Z-scores overexpressed in mutant-GIST determined by LFQ and/or s-SILAC quantitation (FDR<0.05). (**D**) Associated pathways/functions of kinases overexpressed in *KIT*- and *PDGFR*A-mutant GIST vs. normal tissues determined by quantitative MIB-MS profiling. **(E)** Bar graph depicting viability scores for siRNA library screen targeting high-confident kinases elevated in *KIT*- and *PDGFRA*-mutant GIST in GIST-T1+Cas9 and GIST-T1+D842V KIT^KO^ cell lines as measured by Cell Titer Blue assay. siGL2 was negative control, viability score =1.0. Two independent replicates were performed per cell line. **(F)** Quantitative RT-PCR confirmed >70% knockdown of Wee1 (top) and MAP3K3 (bottom) mRNA in both cell lines. Expression levels were normalized to HPRT. *** P < 0.0001; Data represent mean ± SD.

### MK-1775 and avapritinib have enhanced combination effects on *in vitro* GIST cell growth

Although avapritinib has demonstrated dramatic responses in *PDGFRA*-mutant GIST harboring the D842V mutation, acquired resistance to this monotherapy has been observed. A large body of evidence suggests that targeting multiple tumor signaling pathways simultaneously may lead to more sustained tumor control. Given the promising kinome profiling data which demonstrated increased Wee1 activation in GIST (**Figure 3B-C**) and the significant effect on cell viability associated with Wee1 knockdown (**Figure 3E**), we tested the effects of combined inhibition of Wee1 using MK-1775, a commercially available selective inhibitor of Wee1, with KIT/PDGFRA inhibition using avapritinib. We evaluated the effects of MK-1775 and avapritinib using the GIST-T1+Cas9 (KIT driven) and GIST-T1+D842V KIT^KO^ (PDGFRA driven) cell lines, as single agents and in combination at increasing molar ratios. **Figures 4A and B** show single agent dose response curves for GIST-T1+Cas9 and GIST-T1+D842V KIT^KO^, respectively. We first estimated the LD50 for each agent in the two cell lines (**Figures 4A** and **B, left panels**). We then treated each line with increasing doses of the two drugs in a fixed ratio as their LD50s (**Figures 4A** and **B, third panel**). To quantify synergy, combination index (CI) values were calculated (**Figures 4A** and **5B, last panel:** CI values <1 are considered synergistic). The CI_LD50_ values for GIST-T1+Cas9 and GIST-T1+D842V KIT^KO^ were 1.06 and 0.589, respectively, indicating possible synergy only in the PDGFRA-driven cell line, which was then established to be significant via a bootstrap statistic (36). Although synergy was not observed in GIST-T1+Cas9 cell line, a clear additive effect of the two drugs was detected.

**Figure 4.**
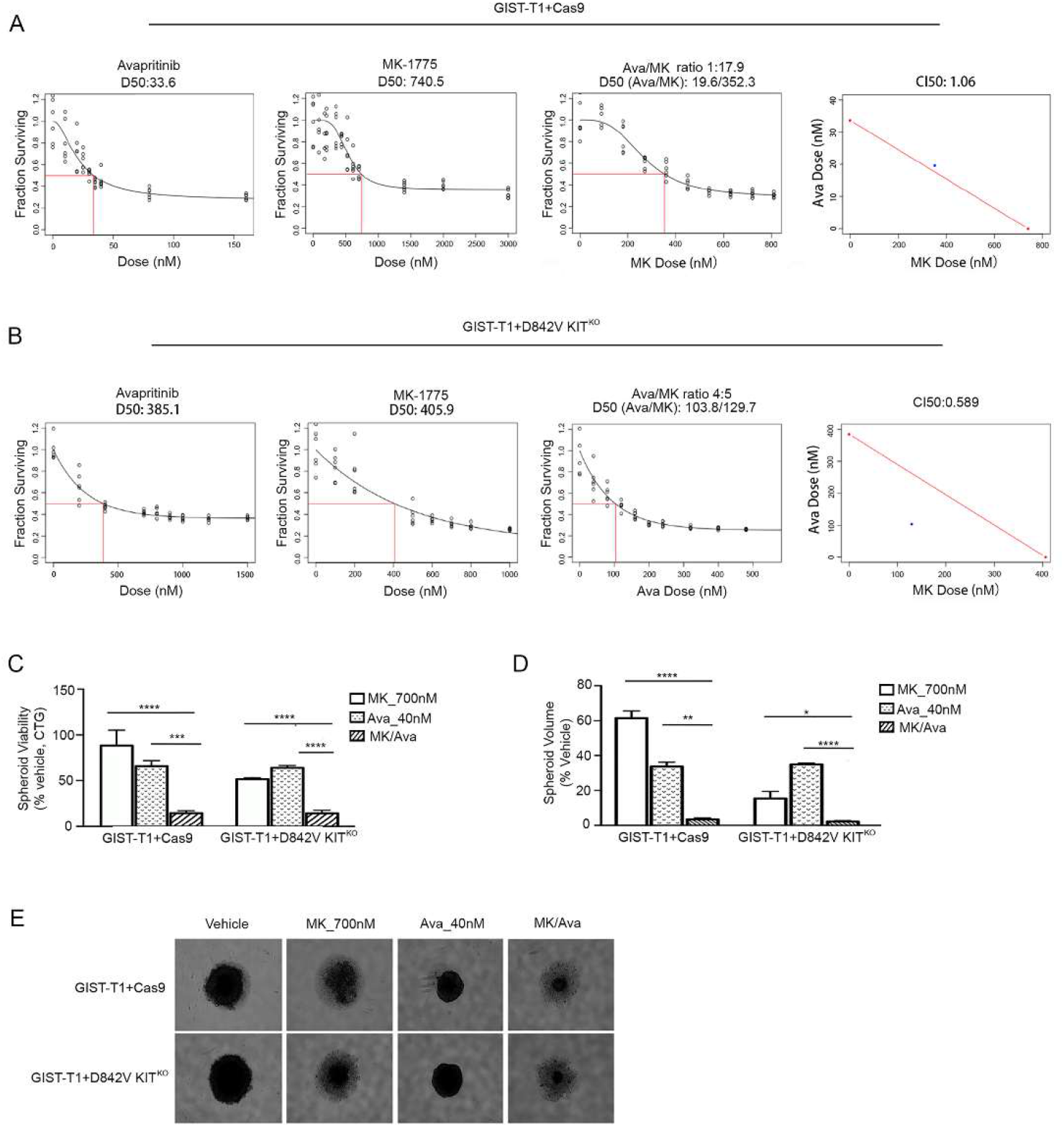
MK-1775 and avapritinib have enhanced combination on *in vitro* GIST cell growth **Panels 1 & 2**, Dose response curves for single agents (avapritinib, MK-1775) in GIST-T1+Cas9. **(A)** and GIST-T1+D842V KIT^KO^ **(B)** cell lines. Red box indicates estimation of LD50 concentration for each single drug. **Panel 3**, Dose response curve representing increasing series of combinations in GIST-T1+Cas9 **(A)** and GIST-T1+D842V KIT^KO^ **(B)** cell lines. Red box indicates estimation of LD50 concentration for combination of drugs. **Panel 4**, Single point (blue) on isobole curve for 50% kill. Red line indicates 50% isobole for strictly additive effect. CI_LD50_ in GIST-T1+Cas9 is 1.06 and not found in the synergistic triangle (region below the red line) **(A)**. CI_LD50_ is 0.589 in GIST-T1+D842V KIT^KO^ and is found within the synergistic triangle **(B)**. Bars represent average viability ± SEM after 120-hour treatment at indicated drug concentrations for GIST-T1+Cas9 and GIST-T1+D842V KIT^KO^ spheroids as a percentage of vehicle-treated spheroids **(C)**. Bars represent the average spheroid volume ± SEM of GIST-T1+Cas9 and GIST-T1+D842V KIT^KO^ spheroids as a percentage of vehicle-treated spheroids **(D)**. All spheroid data were analyzed using GraphPad Prism, with comparisons of treatment groups performed in one-way ANOVA and post hoc comparisons made using Bonferroni multiple comparisons method * p = 0.0165, ** p =0.0046, *** p = 0.0008, **** p =< 0.0001. **(E)** Representative images of GIST-T1+Cas9 and GIST-T1+D842V KIT^KO^ spheroids after 120-hour treatment at indicated concentrations.

To evaluate the effects of the drugs as monotherapies or in combination on 3D GIST cell growth, spheroid assays were performed which more accurately mimic tumor physiology than cells grown in monolayer. GIST-T1+Cas9 and GIST-T1+D842V KIT^KO^ cells form dense, uniformly spherical cultures with true cell-to-cell contacts that are maintained upon physical manipulations, indicative of true speroids (**Figure 4E)**. Treatment of both GIST-T1+Cas9 and GIST-T1+D842V KIT^KO^ spheroids with either of the single agents, MK-1775 (700 nM) or avapritinib (40 nM), resulted in decreased spheroid viability (**Figure 4C)** and volume (**Figures 4D,E)** relative to vehicle-treated spheroids (**Figure 4E)**. However, treatment of these spheroids with the combination resulted in a significantly greater reduction in both viability and volume (**Figures 4C, D, E)**. Interestingly, both MK-1775 as a monotherapy and in combination with avapritinib had greater efficacy in GIST-T1+D842V KIT^KO^ compared to GIST-T1+Cas9 spheroids.

### Combination treatment increases DNA damage and apoptosis

The effect of pharmacological inhibition of KIT/PDGFRA and Wee1 on cell cycle dynamics in GIST cells was measured with a BrdU assay. GIST-T1+Cas9 and GIST-T1+D842V KIT^KO^ cells treated with vehicle, MK-1775, avapritinib or the combination were analyzed by flow cytometry after BrdU incorporation and subsequent antibody binding in combination with direct 7-AAD staining **(Figures 5A, B)**. MK-1775 treatment induced G2-phase cell cycle arrest in GIST-T1+Cas9 cells, whereas cells treated with avapritinib exhibited significant G0/G1-phase arrest compared to control cells. GIST-T1+Cas9 cells treated with the combination exhibited increased subG1 population indicating increased apoptosis compared to either monotherapy treatment group **(Figure 5A)**. Conversely, MK-1775 induced G0/G1-phase arrest in GIST-T1+D842V KIT^KO^ cells whereas avapritinib induced G2 arrest compared to control cells. Combination treatment significantly increased the subG1 population compared to either monotherapy group **(Figure B)**. The subG1 population was two-fold higher in combination-treated GIST-T1+D842V KIT^KO^ cells compared to GIST-T1+Cas9 cells. In order to interrogate the mechanism of action of these inhibitors, we performed immunoblotting on GIST cell lines treated with MK-1775, avapritinib, or the combination **(Figure 5C)**. Following avapritinib treatment, inhibition of KIT and PDGFRA were observed in GIST-T1+Cas9 and GIST-T1+D842V KIT^KO^, respectively. Wee1 typically inhibits CDC2 (cell division cycle protein 2; also known as cyclin dependent kinase 1 (CDK1)) activity by phosphorylating it on two different sites, Tyr15 and Thr14, thereby decreasing its kinase activity and preventing entry into mitosis. Treatment with MK-1775 led to significant inhibition of Tyr15 on CDC2 **(Figure 5C)**. Interestingly, both GIST-T1+Cas9 and GIST-T1+D842V KIT^KO^ cells treated with MK-1775 alone or in combination with avapritinib demonstrated increased γ-H2AX and cleaved-PARP, suggesting increased DNA double-strand breaks and apoptosis. We hypothesize that this increased DNA damage may be a result of loss of cell cycle checkpoints and decreased time for DNA repair mechanisms ultimately causing increased cell death. Interestingly, the KIT-independent cell line, GIST-T1+D842V KIT^KO^, has significantly more cyclin D1 than the KIT-dependent line, GIST-T1+Cas9, in accordance with a recent report (37), providing a potential explanation for the differential effects of MK-1775 and avapritinib in these two cell lines.

**Figure 5.**
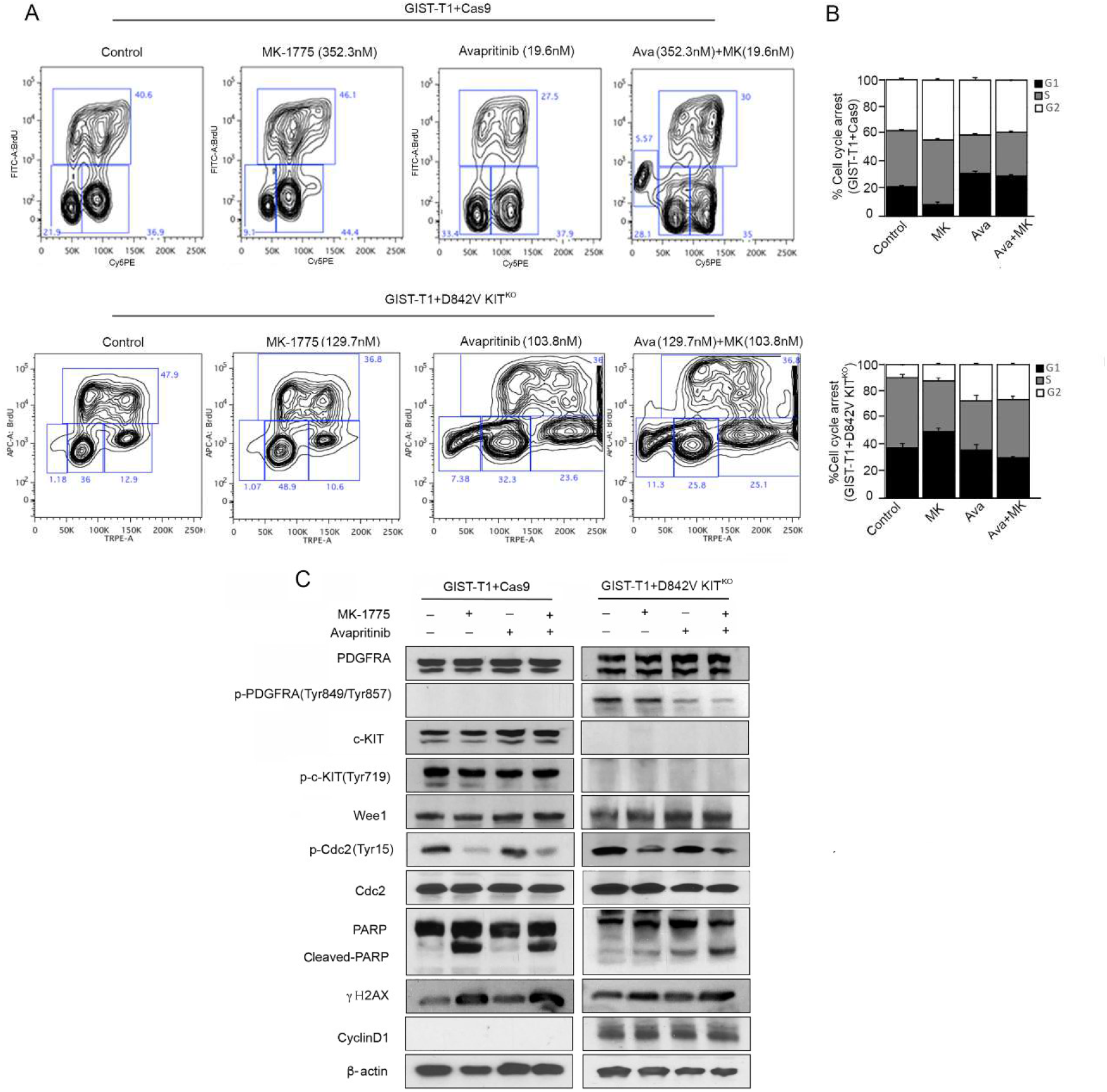
Mechanism of MK-1775 and avapritinib combination in KIT-dependent and – independent GIST cell lines. **(A)** Representative flow cytometry plots and **(B)** quantification of BrdU incorporation in GIST-T1+Cas9 (**upper panel**) treated with 352.3nM MK-1775, 19.6nM Avapritinib and combination for 72h. Statistically significant differences were observed between the following comparisons: for G1 arrest,vehicle vs. avapritinib (p < 0.0009) and vehicle vs. avapritinib/MK-1775 (p < 0.0002); for G2 arrest,vehicle vs. MK-1775 (p < 0.005). **(A)** Representative flow cytometry plots and **(B)** quantification of BrdU incorporation in GIST-T1-D842V+ KIT^KO^ treated **(bottom panel)** with 129.7nM MK-1775, 103.8nM Avaritinib and combination for 72h. Statistically significant differences were observed between the following comparisons: for G1 arrest, vehicle vs. MK-1775 (p < 0.0001); for G2 arrest,vehicle vs. avapritinib (p < 0.005), vehicle vs. avapritinib/MK-1775 (p < 0.0002). Data represent mean ± SD. **(C)** Immunoblot assays of WCEs from GIST-T1+Cas9 (KIT-dependent) and GIST-T1+D842V KIT^KO^ (KIT-independent) cell lines treated as in (**A, B**). Equal concentrations (45-90ug) of WCE from each sample were subjected to immunoblotting with specific antibodies, as indicated. β-actin served as a loading control.

### Combination treatment reduces tumor growth and improves survival *in vivo*

On the basis of these strong *in vitro* data, we hypothesized that there would be benefit in simultaneously inhibiting KIT/PDGFRA and Wee1 leading to loss of cell cycle checkpoint arrest, increased DNA damage and ultimately increased cell death. To test this hypothesis, we performed a GIST xenograft study under a FCCC Institutional Animal Care protocol using the GIST-T1+Cas9 and GIST-T1+D842V KIT^KO^ cell lines. Xenografts were established subcutaneously in a total of 32 C.B17 SCID mice per cell line and randomized into four treatment arms: arm 1: vehicle, arm 2: MK-1775, arm 3: avapritinib and, arm 4: avapritinib/MK-1775 combination. GIST-T1+Cas9 xenografts showed significant disease stabilization in both the avapritinib monotherapy (p = 0.05) and avapritinib/MK-1775 combination (p = 0.002) arms compared to all other groups **(Figure 6A)**. Significant disease stabilization was observed in GIST-T1+D842V KIT^KO^ xenografts in both avapritinib (p = 0.002) and MK-1775 (p = 0.02) monotherapy arms **(Figure 6B)**. Combination treated GIST-T1+D842V KIT^KO^ tumors showed disease stabilization and tumor regression (p = < 0.0002) at day 15 **(Figure 6B)**. Importantly, GIST-T1+Cas9 tumor response led to significant improvement in disease-specific survival in the avapritinib/MK-1775 combination treated group (p = <0.0001) compared to vehicle group **(Figure 6C)**. Kaplan-Meier curves for disease-specific survival of GIST-T1+D842V KIT^KO^ tumors demonstrate that avapritinib/MK-1775 combination-treated mice survived significantly longer than all other mice, including avapritinib alone **(Figure 6D)**. Impressively, at the end of the study (89 days) 75% of the combination treated mice were still alive, one without a measureable tumor, where as no other vehicle and montherapy treated mice were alive. After treatment discontinuation, we observed regrowth of these tumors after approximately four weeks in all but one mouse, whose tumor never regrew.

**Figure 6.**
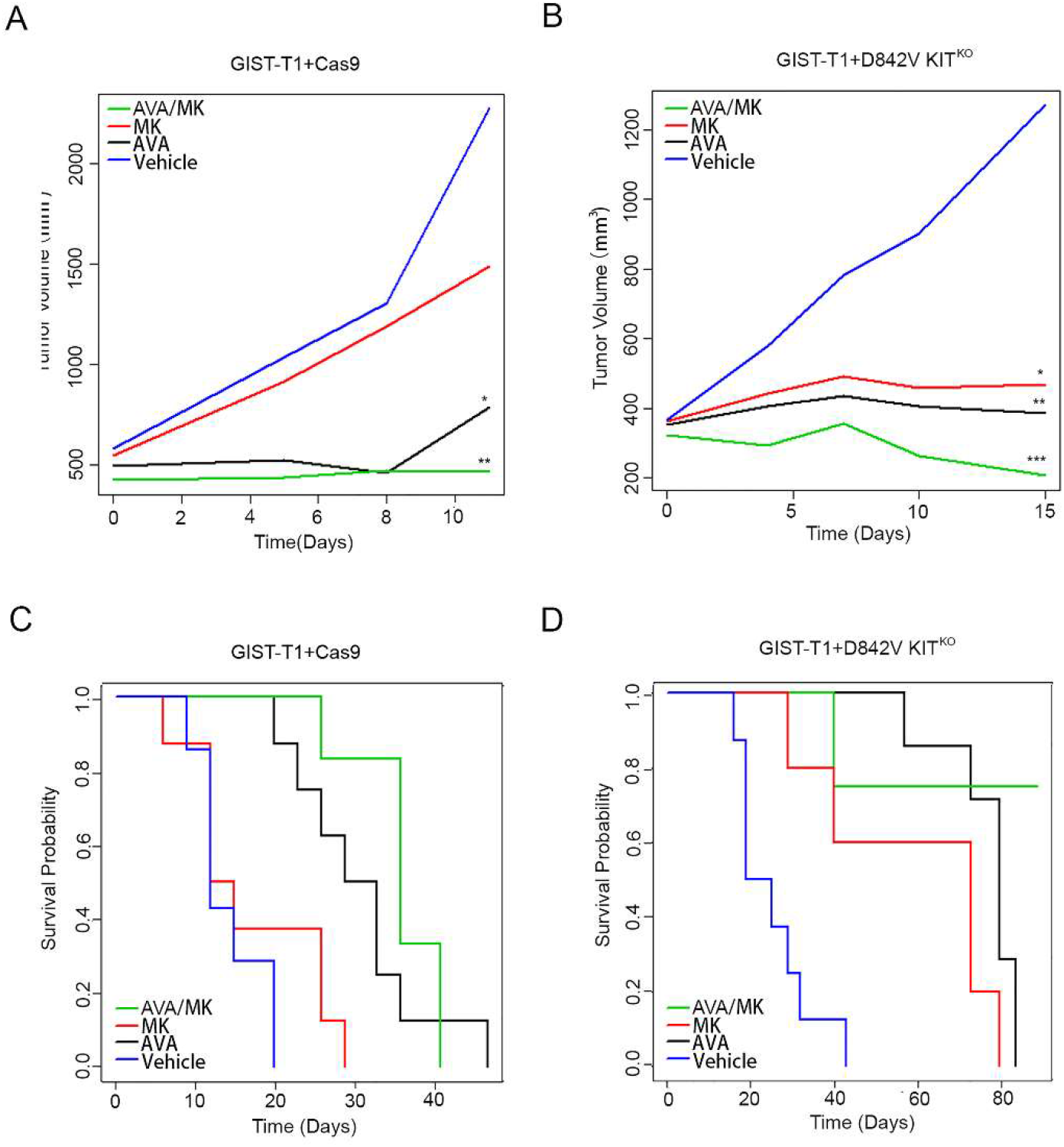
The combination of MK-1775 and avapritinib significantly inhibits GIST growth *in vivo* and improves disease-specific survival. **(A)** Statistically significant decreases in the rate of GIST-T1+Cas9 xenograft tumor growth were observed due to treatment with avapritinib (*p = 0.05, black) and avapritinib+MK-1775 combination (**p = 0.002, green) compared with vehicle group (blue) at day 11. **(B)** Statistically significant decreases in the rate of GIST-T1+D842V KIT^KO^ xenograft tumor growth were observed due to treatment with avapritinib (**p = 0.002) and MK-1775 (* p = 0.02) and avapritinib+MK-1775 (***p = < 0.0002) compared to vehicle group at day 15. Smoothed tumor growth curves (tumor volume vs. time) were computed for each treatment using the lowess smoother in the R statistical language. **(C)** Kaplan-Meier estimate of the probability of disease-specific survival of GIST-T1+Cas9 xenografts. Statistically significant differences (even after adjusting for multiple testing) in disease-specific survival were observed between the following comparisons: Vehicle vs. avapritinib (p< 0.0001), vehicle vs. avapritinib/MK-1775 (p< 0.0001), and MK-1775 vs. avapritinib/MK-1775 (p< 0.0001). **D)** Kaplan-Meier estimate of the probability of disease-specific survival of GIST-T1+D842V KIT^KO^ xenografts. Statistically significant differences (even after adjusting for multiple testing) in disease-specific survival were observed between the following comparisons: Vehicle vs. MK-1775 (p = 0.01), vehicle vs. avapritinib (p < 0.0001), vehicle vs. avapritinib/MK-1775 (p<0.0001), MK-1775 vs. avapritinib/MK-1775 (p =0.01) and avapritinib vs. avapritinib/MK-1775 (p =0.02) are significant. The overall test is also significant (p< 0.0001).

## Discussion

Historically, treatment for advanced GIST involved the sequential application of IM, sunitinib, and regorafenib, regardless of genotype. This approach provided initial benefit to particular molecular subsets of GIST (e.g. *KIT* mutants) and little to no benefit to others (e.g. *PDGFRA* D842V mutants). An increased understanding of GIST biology has revealed clear heterogeneity among the molecular subtypes and a corresponding need for novel therapeutics to target subtype-specific GIST. Recently, this has been borne out with the success of avapritinib in the treatment of *PDGFRA* D842V-mutant GIST, prompting FDA approval of avapritinib as front-line therapy for this subtype in the unresectable or metastatic setting (12). While the application of inhibitors targeting the primary mutant isoforms of KIT and PDGFRA has revolutionized the treatment of GIST, acquired resistance remains a significant clinical challenge. Addressing this challenge may require the identification and targeting of additional protein kinases within cancer-promoting cell signaling pathways that are active within GIST sub-types.

In this study, we utilized a novel chemical proteomics approach, a SILAC-based MIB-MS platform, to profile the kinome of human gastric GIST specimens along with normal gastric tissue. This platform provided a quantitative assessment of kinase abundance for nearly 60% of the human kinome. The kinomes of GIST primary tumors exhibit both a higher level of quantifiable activated kinases, and a distinct profile, as compared to normal gastric tissues. This was not surprising since GIST are generally characterized by gain-of-function mutations that activate multiple signaling pathways. Kinome profiling also revealed significant differences between RTK-mutant (*KIT/PDGFRA)* GIST tumors and SDH-d GIST that lack these mutations. This was expected given the distinct biology of KIT/PDGFRA-driven tumors and SDH-deficient GIST (38, 39). Surprisingly, we also found that *PDGFRA*-mutant GIST tumors expressed a distinct kinome pattern compared to *KIT*-mutant tumors. Interestingly, these differences are partly due to elevated immune cell-associated kinases, including HCK, LCK, BTK, CSF1R and MERTK. Two recent reports (27, 40) have used RNA-seq to obtain immune profiles in GIST. Vitiello *et al*. (27) profiled 75 GIST (N= 37 *KIT*- and 24 *PDGFRA*-mutant) and observed significantly more immune cells present in the *PDGFRA* cohort. While Pantaleo *et al*. (40) did not report genotype specific differences in immune infiltrates in their cohort, their sample size was significantly smaller (N= 21 *KIT*- and 10 *PDGFRA*-mutant), and some of these cases had IM treatment or were classified as unknown treatment status, which could potentially influence the number and activity of immune infiltrates.

Our GIST kinome profiling identified several well-studied and established kinases, such as KIT, PRKCQ (29) and FGFR1 (30), as significantly activated kinases in all GIST compared to normal tissue. In addition, our profiling identified other potential targets. We selected Wee1, gatekeeper of the G2/M cell cycle checkpoint to evaluate, as it was highly abundant in tumors compared to normal tissue and a largely understudied kinase in GIST. Wee1 has been reported to be highly expressed in numerous malignancies including breast, hepatocellular, lung, melanoma and others (33). To assess the role of Wee1, we utilized the Wee1 inhibitor MK-1775 (adavosertib, AZD1775), which has been evaluated in numerous preclinical and clinical trials as single agent or in combination, often with DNA damaging agents (41-43). Notably, recent reports have highlighted significant synergistic potential for MK-1775 in combination with other kinase inhibitors, including TAK228 (34) and alisertib (35). Our loss-of-function studies targeting Wee1 in an isogenic pair of cell lines driven by KIT (GIST-T1+Cas9) or PDGFRA (GIST-T1+D842V KIT^KO^) revealed an essential role for Wee1 in GIST cell proliferation, suggesting Wee1 as a plausible drug target in GIST. We demonstrated enhanced drug combination effects between avapritinib and MK-1775 in both KIT and PDGFRA-driven cell lines using two-dimensional (2D) and three-dimensional (3D) *in vitro* viability studies. Whereas additive effects of the combination were observed in GIST-T1+Cas9 cells, strong synergy was observed in GIST-T1+D842V KIT^KO^ cells treated with the combination. BrdU assays indicated significant differences in the effects of both MK-1775 and avapritinib on cell cycle between the two cell lines, and significantly enhanced apoptosis in the PDGFRA-driven cell line compared to its KIT-driven counterpart. We believe that these differences are due in part to differential expression of cyclin D1, a regulator of the G1/S cell cycle checkpoint, which was recently identified as an oncogenic mediator in KIT-independent GIST (38). Increased expression of γH2AX suggests increased DNA damage, most likely due to loss of cell cycle checkpoint, is responsible for enhanced cell death in combination treated cells.

The results of these *in vitro* studies provided justification for investigating such an approach *in vivo* to determine whether this combination would improve efficacy of avapritinib and/or increase time to resistance in GIST xenografts. Similar to the *in vitro* studies, the avapritinib+MK-1775 combination was significantly better at repressing tumor growth compared to both single agents in both xenograft models, however, tumor regression was only observed in the GIST-T1+D842V KIT^KO^ line. Interestingly, MK-1775 alone had a significant effect on tumor volume compared to vehicle in only the PDGFRA-driven xenografts, indicating inherent cell cycle differences in KIT-versus PDGFRA-driven GIST. These differences were most noticeable when examining disease-specific survival. Impressively, at the end of the study (89 days) in the combination treated arm, 75% of the mice were alive, whereas no other mice, including in avapritinib monotherapy group, survived. Together, these xenograft studies provide strong evidence to support future clinical studies evaluating the use of avapritinib in combination with Wee1 inhibitors in patients with *PDGFRA*-mutant GIST and IM-refractory *KIT*-mutant GIST.

During the preparation of this manuscript, Liu *et al*. (44) published a report examining Wee1 in GIST. They reported elevated expression of Wee1 in GIST compared to normal gastric tissues and a significant anti-proliferative effect of Wee1 knockdown and MK-1775 treatment with DNA damage induction and increased apoptosis. These findings concur with our findings. However, their studies involved *KIT*-mutant GIST only. Our work indicates that in addition to *KIT*-mutant GIST, Wee1 may be a more promising target in *PDGFRA*-mutant GIST. We also hypothesize that Wee1 could be a novel target in SDH-d GIST based on our kinome profiling data and its independence of KIT. Furthermore, we expanded our analysis to include not only *PDGFRA*-mutant GIST cell lines but also to *in vivo* studies of MK-1775, while Liu *et al*. limited studies to *in vitro* evaluations of MK-1775 in *KIT*-mutant GIST. Therefore, our work underscores and expands the evidence for Wee1 serving an important role in GIST biology, and provides a strong rationale for the therapeutic targeting of Wee1 in all subtypes of GIST.

## Material and Methods

### Kinome Profiling Experimental Design, Data Analysis and Statistical Rationale-

For proteomic measurement of kinase abundance in tissues, we used MIB-MS profiling and quantitated kinase levels using a combination of Label-Free Quantitation (LFQ) and super-SILAC (s-SILAC) (22, 45). Briefly, an equal amount of s-SILAC reference (5 mg) was spiked into each primary tissue sample (5 mg) and kinases purified from tissues using MIB-resins, kinases were eluted, digested, and peptides analyzed by LC-MS/MS as previously described (19). To identify differences in kinase abundance amongst GIST tissues, we performed MIB-MS analysis on GIST (n=33) (*KIT*-mutant (n=15), *PDGFRA*-mutant (n=10), SDH-d (n= 8)) and normal gastric tissues (n=9). Measurement of MIB-enriched kinase abundance in tissues was performed by LFQ and s-SILAC quantitation using MaxQuant software version 1.6.1.0.

### Data analysis of MIB-MS

MaxQuant normalized LFQ values or SILAC ratios (H/L) were filtered for human protein kinases in excel and then imported into Perseus software (1.6.2.3) for quantitation. <u>LFQ data processing:</u> Kinase LFQ values were filtered in the following manner: kinases identified by site only were removed, reverse or potential contaminants were removed then filtered for kinases identified by >1 unique peptide. Kinase LFQ intensity values were then log2 transformed, technical replicates averaged, and rows filtered for minimum valid kinases measured (n= >70% of runs). No imputation of missing values was performed. Filtered LFQ data was annotated and subjected to a Student’s *t*-test comparing GIST tissue subtypes using Perseus Software. Parameters for the Student’s *t*-test were the following: S0=0.1, side both using Permutation-based FDR <0.05. <u>s-SILAC data processing:</u> Kinase s-SILAC ratios were transformed 1/(x) to generate light / heavy ratios, log2 transformed, technical replicates averaged, and rows filtered for minimum valid kinases measured (n= >70% of runs). No imputation of missing values was performed. Filtered normalized s-SILAC ratios were annotated and subjected to a Student’s t-test comparing GIST tissue subtypes using Perseus Software. Parameters for the Student’s *t*-test were the following: S0=0.1, side both using Permutation-based FDR <0.05. Volcano plots depicting differences in kinases abundance were generated using R studio software. For PCA analysis of kinase log2 LFQ values, rows were filtered for kinases measured in 100% of MIB-MS runs and principal component analysis (PC1 vs PC2, PC2 vs PC3 and PC1 vs PC3) performed to visualize kinome profiles amongst tissue samples. Scatterplots or bar graphs were used to compare LFQ *vs*. s-SILAC measurements of differentially expressed kinases amongst tumor and normal tissues. Plots comparing differences in kinase log2 LFQ values or kinase log2 s-SILAC ratios were determined by Student’s *t*-test. Scatter plots depicting differences in kinases abundance were generated using R studio software and bar graphs generated in excel or Prism.

### Nano-LC-MS/MS

Proteolytic peptides were resuspended in 0.1% formic acid and separated with a Thermo Scientific RSLC Ultimate 3000 on a Thermo Scientific Easy-Spray C18 PepMap 75µm x 50cm C-18 2 μm column. For MIB runs, a 240 min gradient of 4-25% acetonitrile with 0.1% formic acid was used. For total proteome runs, a 305 min gradient of 2-20% (180 min) 20%-28% (45 min) 28%-48% (20 min) acetonitrile with 0.1% formic acid was used. Both gradients were run at 300 nL/min at 50°C. Eluted peptides were analyzed by Thermo Scientific Q Exactive or Q Exactive plus mass spectrometer utilizing a top 15 methodology in which the 15 most intense peptide precursor ions were subjected to fragmentation. The AGC for MS1 was set to 3×10^6^ with a max injection time of 120 ms, the AGC for MS2 ions was set to 1×10^5^ with a max injection time of 150 ms, and the dynamic exclusion was set to 90 s.

### Proteomics Data Processing

Raw data analysis of LFQ or s-SILAC experiments was performed using MaxQuant software 1.6.1.0 and searched using Andromeda 1.5.6.0 against the Swiss-Prot human protein database (downloaded on April 24, 2019, 20402 entries). The search was set up for full tryptic peptides with a maximum of two missed cleavage sites. All settings were default and searched using acetylation of protein N-terminus and oxidized methionine as variable modifications. Carbamidomethylation of cysteine was set as fixed modification. The precursor mass tolerance threshold was set at 10 ppm and maximum fragment mass error was 0.02 Da. LFQ quantitation was performed using MaxQuant with the following parameters; LFQ minimum ratio count: 2, Fast LFQ: selected, LFQ minimum number of neighbors; 3, LFQ average number of neighbors: 6. SILAC quantification was performed using MaxQuant by choosing multiplicity as 2 in group-specific parameters and Arg10 and Lys8 as heavy labels. Global parameters for protein quantitation were as follows: label minimum ratio count: 1, peptides used for quantitation: unique, only use modified proteins selected and with normalized average ratio estimation selected. Match between runs was employed for LFQ and s-SILAC quantitation and the significance threshold of the ion score was calculated based on a FDR of < 1%.

### MIBs preparation and chromatography

Experiments using MIB/MS were performed as previously described (*25*). Briefly, cells or tumors were lysed and an equal amount of the s-SILAC reference (5 mg) lysate was added to non-labeled (5 mg) lysate (cell, or tumor tissue) and endogenous kinases isolated by flowing lysates over kinase inhibitor-conjugated Sepharose beads (purvalanol B, VI16832, PP58 and CTx-0294885 beads) in 10 ml gravity-flow columns. Eluted kinases were reduced by incubation with 5 mM DTT at 65ධC for 25 minutes, alkylated with 20 mM iodoacetamide at room temperature for 30 minutes in the dark, and alkylation was quenched with DTT for 10 minutes, flowed by digested with sequencing-grade modified trypsin (Promega) overnight at 37ධC. C-18 purified peptides were dried in a speed vac, and subsequent LC-/MS/MS analysis was performed.

### Data Availability

All mass spectrometry proteomics data have been deposited to ProteomeXchange Consortium via PRIDE partner repository with the dataset identifier PXD020720.

### Cell lines, compounds, and antibodies

GIST-T1 tumor cell line possessing a heterozygous mutation in *KIT* exon 11 was kindly provided by Takahiro Taguchi (Kochi University, Kochi, Japan) (46). GIST-T1+Cas9 and GIST-T1+D842V KIT^KO^ are sublines of GIST-T1. GIST-T1+Cas9 was generated transduction of Cas9 using the LentiV-Cas9-puro vector system generously provided by Christopher Vakoc (Cold Spring Harbor Laboratory). GIST-T1+D842V KIT^KO^ subline was created by transducing cells with D842V-mutant PDGFRA. Endogenous KIT expression was knocked-out using CRISPR/Cas9. Knockout was verified at protein and DNA levels. All GIST-T1 cell lines were grown in Iscove’s Modified Dulbecco’s Media (IMDM) with 15% FBS and were routinely monitored by Sanger sequencing to confirm *KIT*/*PDGFRA* mutation status and cell identity. Cell lines were regularly tested for mycoplasma contamination by PCR and MycoAlert Mycoplasma Detection Kit (Lonza). Avapritinib and MK-1775 were obtained from Selleckchem (Houston, TX). For *in vitro* experiments, avapritinib and MK-1775 were dissolved in DMSO. For *in vivo* experiments, avapritinib and MK-1775 were dissolved in 2% DMSO + 40% PEG400 + 2% Tween80 + ddH_2_O. All antibodies used were purchased from Cell Signaling Technology, except β-actin and γ-H2AX (Millipore Sigma, St. Louis, MO).

### siRNA transfection

The custom siRNA library was synthesized with four independent siRNAs pooled per target (siGenome SMARTpool, Dharmacon, Lafayette, CO). Transfection conditions were determined for GIST-T1+Cas9 and GIST-T1+D842V KIT^KO^ cells using siRNA SMARTpools against KIT, PDGFRA and GL-2 (Dharmacon) controls to achieve Z′ factor of 0.5 or greater. Reverse transfection mixtures were assembled in 96-well with final siRNA concentration of 50nM. After seventy-two hours, plates were assayed for cell viability using the CellTiter Blue (CTB) Viability Assay (Promega, Madison, WI) as previously described (47).

### Cell proliferation/viability assay

Tumor cells were plated in 96-well plates at optimal 0.6 ×10^4^ densities and incubated overnight. Wells were treated in sextuplicate with varying doses of MK-1775 and/or avapritinib. Cell proliferation and viability were measured at 72 hours after treatment using CTB Assay as described above. Assays were performed as three independent biological replicates, with minimum of three technical replicates in each treatment arm. An increasing dose series was used for each drug to estimate LD50. A function of form A + (1-A) * exp(-B*dose) or A + (1-A)/[1 + (dose/B)^p] was fitted to data by least squares. These functions were used to interpolate surviving fractions between those in dose series and set to ½ to estimate corresponding LD50s, (LD50-1 and LD50-2). Increasing series of combination doses in same ratio were used as their LD50s to estimate LD50 of that combination (dose1 and dose2), using an interpolating function. If the combination index, CI = dose1/LD50-1 + dose2/LD50-2 < 1 then the point (dose1, dose2) may be synergistic, otherwise, it was considered either additive or antagonistic. If not considered additive or antagonistic, a bootstrap resampling method was used to test the null hypothesis of no synergism (36).

### Spheroid drug sensitivity

Spheroids were formed in 96 Well U-Bottom Clear Cell Repellent Surface Microplates (Greiner Bio-One). GIST-T1+Cas9 and GIST-T1+D842V KIT^KO^ cells were suspended in complete IMDM (4,500 cells/well) for 24 hours for spheroid formation. Spheroids were treated with appropriate drug(s) and were imaged at 4× magnification by EVOS FL Digital Inverted Microscope after 120 hours of treatment. Spheroid surface area and viability were measured and statistical analyses were conducted as described previously (47). Three independent biological replicate experiments were performed with minimum of three technical replicates in each treatment arm.

### BrdU incorporation assay

DNA synthesis proliferation rate was measured using BrdU Flow kit (BD Biosciences, San Jose, CA) according to manufacturer’s protocol. Treated GIST-T1+Cas9 and GIST-T1-D842V+KIT^KO^ cells were labelled with bromodeoxyuridine (BrdU) for 3.5 h. Anti-FITC-BrdU antibody was used in GIST-T1+Cas9 cells and anti-APC-BrdU was used in GIST-T1-D842V+KIT^KO^ cells. Total DNA was stained with 7-amino-actinomycin D (7-AAD). Double labeled samples were analyzed using two-color flow cytometric analysis conducted on LSR ll BD flow cytometer. Data was analyzed and displayed using FlowJo software.

### Preparation of whole cell extract from cells and immunoblot assays

The whole cell extracts were prepared and evaluated by immunoblot assay as described previously (48).

### GIST xenografts and drug administration

All studies involving animals followed procedures approved by the FCCC Institutional Animal Care and Use Committee. GIST-T1+Cas9 and GIST-T1+D842V KIT^KO^ cells were washed and resuspended in PBS at a density of 1× 10^6^ cells/100μL. One hundred microliters of cells in PBS were mixed thoroughly with 100μL of Matrigel Matrix (BD Biosciences) and suspension was injected subcutaneously into the right flanks of SCID mice (CB.17/SCID, Taconic Biosciences). A total of 64 mice were used in this study. Tumor volume was calculated as described previously (47). When tumors reached approximately 300 mm^3^, mice were randomized into five treatment arms: arm 1, vehicle; 2, MK-1775 at 60 mg/kg, twice a day (oral); arm 3 avapritinib 10 mg/kg, once a day (oral) and arm 4, combination of MK-1775 and avapritinib at monotherapy doses. Treatment was continued until tumors exceeded >10% of their body weight or animals demonstrated distress or weight loss >10%.

### Tumor growth modeling

Tumor volume was measured for every mouse in all treatment arms (vehicle, MK-1775, avapritinib and combination) at a total of 15 distinct time points in GIST-T1+Cas9 xenografts, from baseline (Day 0) until study conclusion (47 Days) and 25 distinct time points in GIST-T1+D842V KIT^KO^ xenografts, from baseline (Day 0) until study conclusion (Day 89). A longitudinal model based on generalized estimating equations approach (Gaussian model with identity link and autoregressive correlation structure) was used to model treatment effect and time on (the logarithm of) tumor volume. A linear time-effect was included in the model for logarithm of tumor volume and interacted with treatment. Disease specific survival and tumor volume were compared between treatment groups using log rank and Mann–Whitney tests, respectively. All tests were two-sided and used a type I error of 5%. The package geepack and survival in R statistical language and environment was used in these computations.

## Acknowledgements

This work was supported by NIH CORE grant CA06927 (FCCC), NCI R00 CA158065 (L.R), NCI R01 CA212662 (L.R.), NCI R01 CA211670 (J.S.D), NCI R50 CA211479 (M.B.E), WJ Avery Fellowship (S.Y.) and US Department of Veterans Affairs I01 BX000338 (M.C.H). We would like to acknowledge the following facilities at FCCC for their work contributing to this manuscript: Biosample Repository, Cell Sorting, High Throughput Screening, Genotyping and Real-Time PCR, and the Laboratory Animal Facility. The authors would especially like to thank the David Foundation and the GIST Cancer Research Fund for their continued support.

## Author contributions

SY, LR, JSD, and MVM conceptualized the research, designed the experiments, and wrote the manuscript. SY, DS, MK, JD, MGB and MBE performed experiments and analyzed data. KJL, KD, YD, SL performed statistical analyses. LK and MCH provided *KIT* and PDGFRA-mutant GIST cell lines. RDM provided GIST samples.

---

**Supplemental Figure 1.**
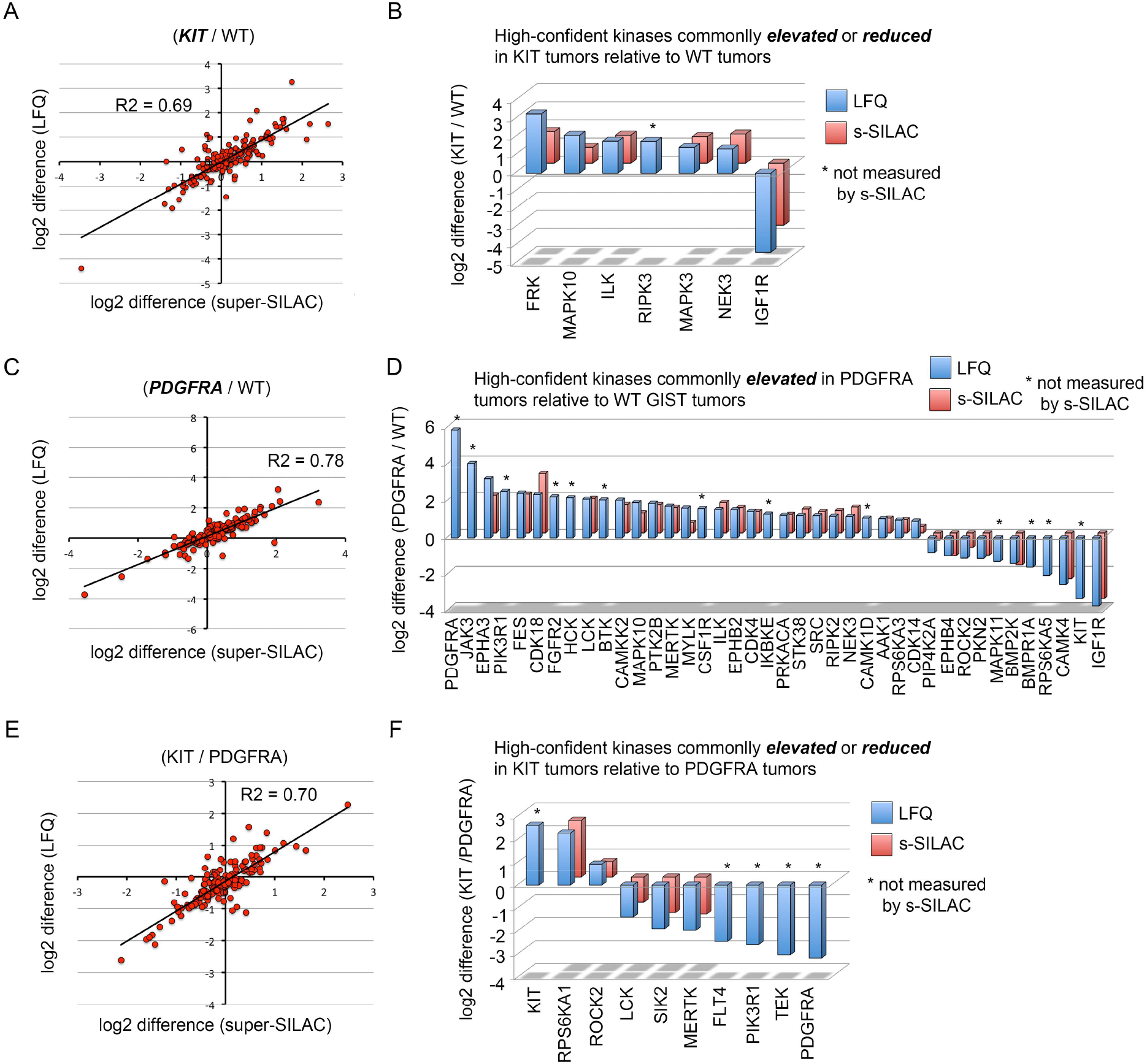
Defining kinome signatures of GIST tumor subtypes using MIB-MS profiling. (**A, C** and **E**) Scatterplot depicts the overlap in kinases elevated or reduced determined by LFQ or s-SILAC comparing (A) *KIT*-mutant vs WT, (C) *PDGFRA*-mutant vs WT and (E) *KIT*-mutant vs. *PDGFRA*-mutant. Regression analysis (R^2^) among quantitative methods was performed in Perseus Software. Differential expressed kinases commonly identified by LFQ and s-SILAC quantitation (FDR <0.05) are labeled. (**B, D** and **F**) Bar graph depicts high-confident kinases log2 LFQ values overexpressed in (B) *KIT*-mutant vs. WT, (D) *PDGFRA*-mutant vs. WT or (F) *KIT*-mutant vs *PDGFRA*-mutant GIST primary tumors determined by LFQ and/or s-SILAC quantitation (FDR <0.05).

